# Benchmarking and optimizing Perturb-seq in differentiating human pluripotent stem cells

**DOI:** 10.1101/2025.01.21.633969

**Authors:** Sushama Sivakumar, Yihan Wang, Sean C Goetsch, Vrushali Pandit, Lei Wang, Huan Zhao, Anjana Sundarrajan, Daniel Armendariz, Chikara Takeuchi, Mpathi Nzima, Wei-Chen Chen, Ashley E Dederich, Lauretta El Hayek, Taosha Gao, Renad Ghazawi, Ashlesha Gogate, Kiran Kaur, Hyung Bum Kim, Melissa K McCoy, Hanspeter Niederstrasser, Seiya Oura, Carolos A Pinzon-Arteaga, Menaka Sanghvi, Daniel A Schmitz, Leqian Yu, Yanfeng Zhang, Qinbo Zhou, W. Lee Kraus, Lin Xu, Jun Wu, Bruce A Posner, Maria H Chahrour, Gary C Hon, Nikhil V Munshi

**Affiliations:** Department of Internal Medicine, Division of Cardiology, University of Texas Southwestern Medical Center, Dallas, TX, USA; Cecil H. and Ida Green Center for Reproductive Biology Sciences, University of Texas Southwestern Medical Center, Dallas, TX, USA; Department of Neuroscience, University of Texas Southwestern Medical Center, Dallas, TX, USA; Eugene McDermott Center for Human Growth and Development, University of Texas Southwestern Medical Center, Dallas, TX, USA; Department of Biochemistry, University of Texas Southwestern Medical Center, Dallas, TX, USA; Department of Molecular Biology, University of Texas Southwestern Medical Center, Dallas, TX, USA; Quantitative Biomedical Research Center, Peter O’Donnell Jr School of Public Health, University of Texas Southwestern Medical Center, Dallas, TX, USA; Department of Pediatrics, Division of Hematology/Oncology, University of Texas Southwestern Medical Center, Dallas, TX, USA; Department of Psychiatry, University of Texas Southwestern Medical Center, Dallas, TX, USA; Center for the Genetics of Host Defense, University of Texas Southwestern Medical Center, Dallas, TX, USA; Peter O’Donnell Jr Brain Institute, University of Texas Southwestern Medical Center, Dallas, TX, USA; Lyda Hill Department of Bioinformatics, Department of Obstetrics and Gynecology, University of Texas Southwestern Medical Center, Dallas, TX, USA; Hamon Center for Regenerative Science and Medicine, University of Texas Southwestern Medical Center, Dallas, TX, USA

## Abstract

Perturb-seq is a powerful approach to systematically assess how genes and enhancers impact the molecular and cellular pathways of development and disease. However, technical challenges have limited its application in stem cell-based systems. Here, we benchmarked Perturb-seq across multiple CRISPRi modalities, on diverse genomic targets, in multiple human pluripotent stem cells, during directed differentiation to multiple lineages, and across multiple sgRNA delivery systems. To ensure cost-effective production of large-scale Perturb-seq datasets as part of the Impact of Genomic Variants on Function (IGVF) consortium, our optimized protocol dynamically assesses experiment quality across the weeks-long procedure. Our analysis of 1,996,260 sequenced cells across benchmarking datasets reveals shared regulatory networks linking disease-associated enhancers and genes with downstream targets during cardiomyocyte differentiation. This study establishes open tools and resources for interrogating genome function during stem cell differentiation.

## INTRODUCTION

Perturb-seq is a high throughput phenotypic assay that combines CRISPR interference (CRISPRi) using a library of single guide RNAs (sgRNAs) with single cell RNA-seq (scRNA-seq) to analyze the effects of thousands of genetic perturbations in a single experiment^1,2^. Perturb-seq studies have identified genes required for survival and cell death, pinpointed regulators of specific genes, and highlighted genes implicated in T-cell exhaustion^2–6^. However, large-scale Perturb-seq experiments have largely been conducted either in established cell lines or post-mitotic primary cells^2,4,7^. While extending Perturb-seq to hPSC (human pluripotent stem cell) differentiation systems would yield insights on development and disease, it is challenging for several reasons. First, hPSCs are prone to transgene silencing and variegated expression, which presents a substantial barrier to stable dCas9-KRAB expression. Second, hPSCs stably repress large segments of their genome during differentiation in a cell type-specific manner^8^, which can impact expression of CRISPRi effectors and sgRNAs. Third, differentiation systems represent a continuum of cell states, making their perturbation and analysis more challenging than established cell lines (e.g. K562). Thus, we aimed to establish an hPSC system that enables stable repression throughout directed differentiation for comprehensive Perturb-seq studies.

Differentiation of hPSCs provides a scalable platform to interrogate the functions of genes and regulatory elements in potentially any human cell type at developmentally relevant time points. For example, Perturb-seq screens can be used to establish the function of individual genes during human cardiomyocyte (CM) differentiation and cardiac development^9–11^. However, so far none of the CRISPRi screens was conducted using dCas9-KRAB knocked into a genomic safe harbor (GSH) locus to ensure stable, long-term transgene expression. Here, we benchmark and optimize experimental procedures to efficiently and cost-effectively perform Perturb-seq during hPSC differentiation into cardiomyocytes and neurons. First, we generated stably integrated CRISPRi machinery in multiple human pluripotent stem cell lines (H9 embryonic stem cells (ESCs) and WTC11 induced pluripotent stem cells (iPSCs)) to ensure robust expression of dCas9-KRAB. Second, we designed sgRNA architectures to ensure on-target repression and efficient detection after scRNA-Seq. Third, we compared 3 multiplexing strategies (lentivirus, piggyBac, recombinase) for sgRNA delivery into hPSCs. Fourth, we developed quality control steps during cardiomyocyte differentiation to ensure optimal library coverage and efficient differentiation into cardiomyocytes. Fifth, we optimized super loading of cells during library preparation of scRNA-seq to maximize cell recovery of transcriptomic data after sequencing. Our comparative analyses inferred the functions of targeted genes and enhancers in cardiac development, and constructed gene regulatory networks linking genetic elements and key developmental genes. Overall, our method provides a rich resource for the scientific community and provides tools and recommendations to build gene regulatory networks that govern early cell development. These best practices represent our standard operating procedures (SOP) for Perturb-seq experiments performed as part of the IGVF consortium^12^.

## RESULTS

### Engineering versatile pluripotent stem cells for diverse Perturb-seq applications

To successfully perform Perturb-seq experiments during differentiation, a robust repression machinery installed in pluripotent cells is required. We engineered pluripotent stem cell lines to stably express the CRISPRi machinery with dCas9 fused with the transcriptional repressor KRAB. Previously, we used PiggyBac transposition to insert dCas9-KRAB into the genome^9^, but the random integration of multiple copies and the potential for transgene silencing over time were problematic. To circumvent these issues, recent studies demonstrate that stably integrated dCas9-KRAB into the CLYBL safe harbor locus of WTC11 iPSCs effectively represses target genes in human differentiated neuronal cells^13,14^. We adapted this approach to expand CRISPRi Perturb-seq to diverse cell types (**Figure 1A**). First, we generated H9 ESCs constitutively expressing heterozygous dCas9-KRAB in the CLYBL locus (H9 dCK) (**Figure 1A, S1A-S1C**), as well as WTC11 iPSCs with heterozygous dCas9-KRAB expression (WTC11 dCK) (**Figure 1A, S1A-C**). To gain temporal control of CRISPRi and to assess genomic function at distinct times during differentiation, we also engineered a DOX-inducible^15^ dCas9-KRAB in H9 ESCs (H9 idCK) by separately installing the rtTA activator cassette into the ROSA26 locus and the TRE-dCas9-KRAB into the CLYBL locus (**Figure 1A, S1A-E**)^16^. This dual targeting approach allows independent transcription of transgenes without interference^15^. Genotyping of all 3 engineered CRISPRi hPSCs validated correct transgene integration (**Figure S1B-E**).

**Figure 1:**
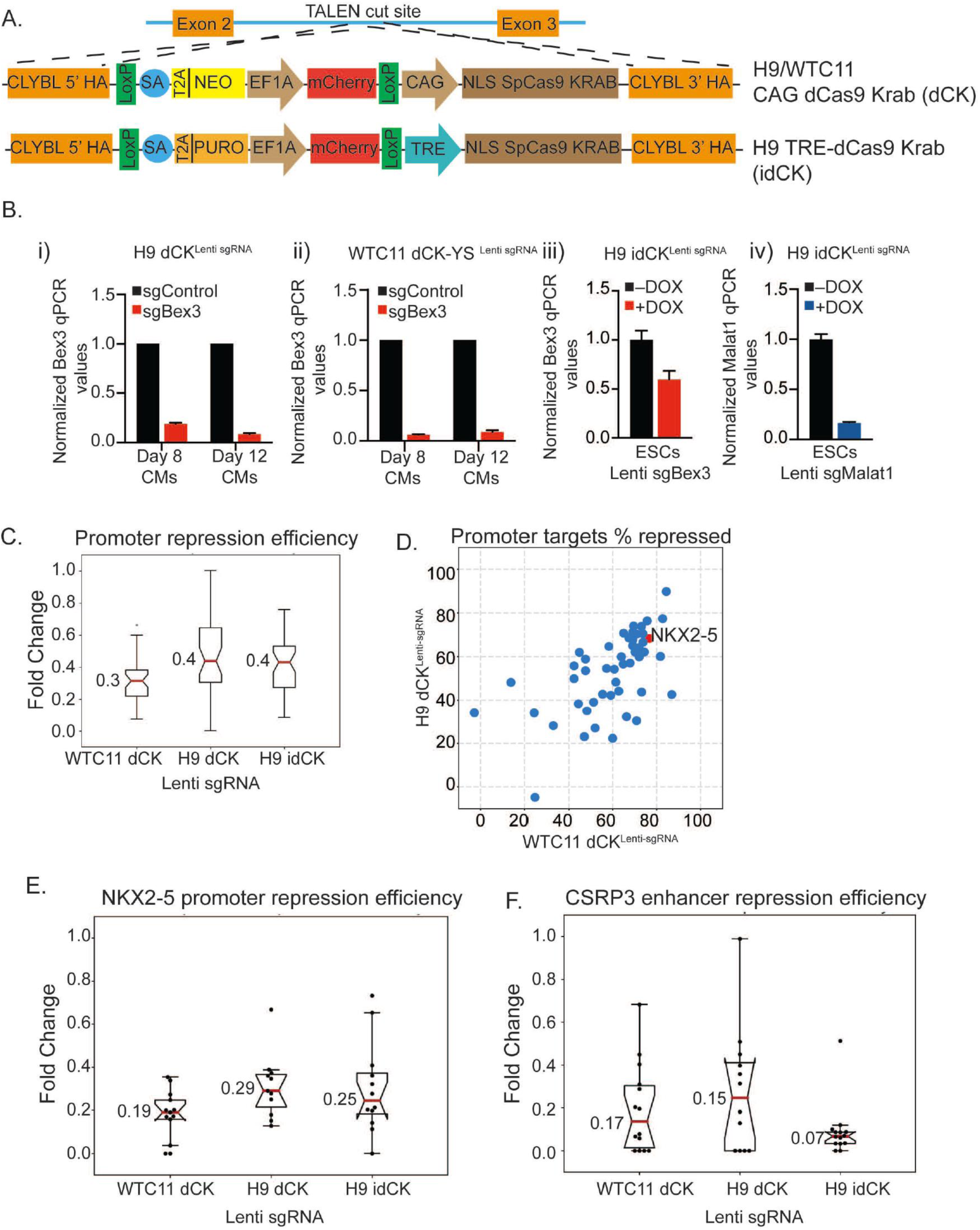
Engineering PSCs for diverse Perturb-seq applications: A) Schematic of transgene integrated in the CLYBL GSH locus in constitutive H9/WTC11 dCK and inducible H9 idCK PSCs. EF1-mcherry selection cassette is placed upstream of dCas9-KRAB to allow selection of cells during ESC line construction. B) qPCR showing repression of indicated targets using lentiviral sgRNAs in (i) H9 dCK, (ii) WTC11 dCK-YS, (iii & iv) H9 idCK C). Graph showing median repression efficiency of all promoters in indicated PSCs infected with lentiviral sgRNAs. D) Correlation map of promoter repression efficiency across H9 dCK and WTC11 dCK. Red target indicates repression of NKX2-5. E) Graph showing median repression efficiency of NKX2-5 promoter in indicated cell lines. Each dot represents a single sgRNA targeting NKX2-5 promoter. F) Graph showing median repression efficiency of CSRP3 enhancer in indicated cell lines. Each dot represents a single sgRNA.

To compare the performance of constitutive and inducible CRISPRi stem cells, we introduced sgRNAs using a lentiviral strategy (**Figure 1B**). Bulk qPCR experiments confirmed efficient repression (80-95%) of positive control sgRNAs targeting BEX3, MALAT1, or TFRC^9^ in PSCs and during CM differentiation (**Figure 1B**). We tested two independently derived WTC11 lines: WTC11 dCK derived from an independent study (WTC11-dCK-YS)^14^ and WTC11 dCK engineered in-house (WTC11-dCK-NVM). Both performed equally at repressing targets (**Figure 1B(ii), S1F**). Using H9 idCK, we obtained ∼50% repression of lentiviral sgBEX3 and ∼80% repression of lentiviral sgMALAT1 after 48h of DOX treatment to turn on dCas9 expression **(Figure 1B(iii-iv)**). Longer treatment may allow better repression, particularly for genes with higher transcript levels and/or longer half-lives.

To comprehensively compare the engineered lines, we designed an sgRNA library targeting 193 cardiac promoters and enhancers (Cardiac pilot; IGVF data portal), cloned into a vector containing an EF1a-mTagBFP-puro selection cassette for dynamic monitoring of lentiviral expression and estimation of multiplicity of infection (MOI). After lentiviral infection of stem cells at low MOI (1 sgRNA/cell) and sorting for BFP+ cells, we differentiated to cardiomyocytes and performed Perturb-seq. Examining promoters, we observed robust CRISPRi repression in WTC11 cells (∼70% knockdown efficiency), H9 cells (∼60% knockdown efficiency), and inducible H9 cells (∼60% knockdown efficiency) (**Figure 1C**). We also observed a strong correlation in promoter knockdown efficiencies across WTC11 and H9 cells (**Figure 1D**), indicating consistent sgRNA mediated promoter repression across lines. Repression of well-known cardiac transcription factor NKX2-5 by CRISPRi resulted in ∼70-80% decrease in transcript levels across cell lines as measured by scRNA-seq (**Figure 1E**). Next, we measured the impact of CRISPRi repression of enhancers on proximal target genes (distance <1Mbp) and observed reproducible knockdown for strong enhancers across all engineered cell lines. For example, CSRP3 enhancer repression strongly decreased CSRP3 expression (**Figure 1F**), and repression of an IRX4 enhancer led to a decrease in IRX4 mRNA levels by 80-90% in both H9 dCK and WTC11 dCK PSCs (**Figure S1G**). In summary, these observations of strong CRISPRi repression efficiency across our engineered cell lines and across diverse genomic targets support the use of this toolkit in future Perturb-seq studies to examine genome function (**Figure S1H**).

### Comparing sgRNA delivery methods to optimize target gene repression

An important parameter for Perturb-seq is effective delivery and detection of sgRNAs during single-cell RNA-Seq. In addition to the lentiviral strategy described above, we also tested sgRNA delivery by PiggyBac (PB) transposition and PA01 recombinase integration (**Figure 2A**). A summary of the advantages and disadvantages of each delivery method are listed in **Supplementary Table 1**. We designed the PB vector based on the same architecture as the successful lentivirus vector, with a U6-sgRNA cassette upstream of the EF1-mTagBFP-puro selection cassette (**Figure 2B**). In bulk experiments with control sgRNAs delivered by PB, we observed 80-90% repression of target genes in both H9 dCK and WTC11 dCK lines (**Figure 2C, S2A-D**), with weaker repression in the inducible H9 idCK cells (**Figure 2C(iii)**). The in-house generated WTC11-dCK-NVM was also efficient at repressing PB-sgRNAs against BEX3 and TFRC targets (**Figure S2C-D**). Given the consistency between lines, we chose to use the WTC11-dCK-YS line^14^ for CM differentiation Perturb-seq experiments.

**Figure 2:**
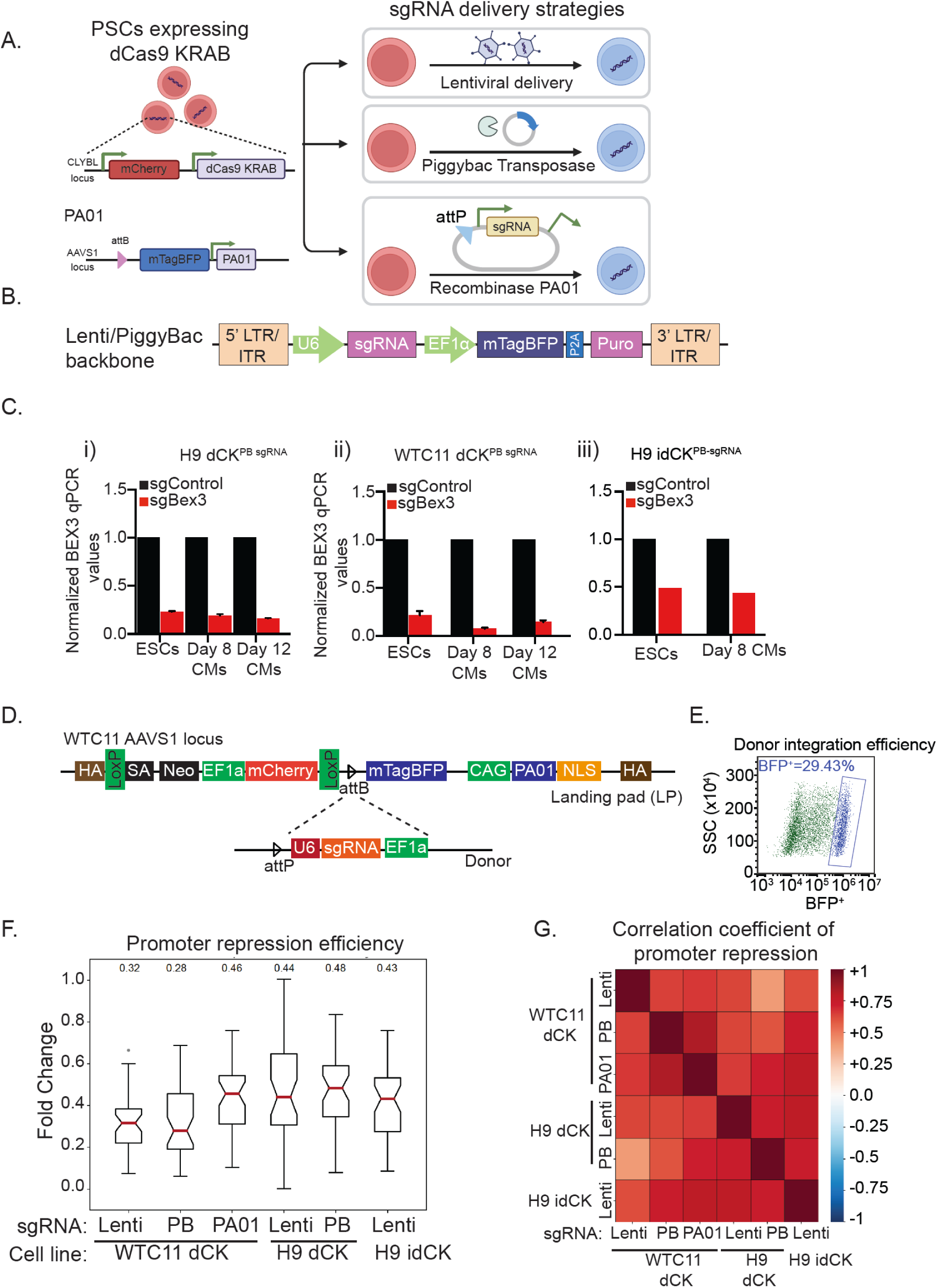
Comparing sgRNA delivery methods to optimize target gene repression: A) Schematic of lentiviral (LV) or Piggybac (PB)-transposon or recombinase PA01 mediated sgRNA delivery into dCK PSCs (WTC11/ H9). B) schematic of U6-sgRNA construct that is nucleofected (PB) or used to infect (LV) cells. Placing the U6-sgRNA cistron upstream of the EF1-mTagBFP-puromycin selection cassette allows for optimum expression of U6-sgRNA, efficient capture of sgRNAs during sc-RNAseq and good target repression. C) qPCR showing repression of Bex3 target in i) H9 dCK, ii) WTC11 dCK and iii) H9 idCK PSCs or differentiated CMs nucleofected with PB Bex3 sgRNA or non-targeting sgRNA. D) Schematic of recombinase PA01 landing pad inserted into AAVS1 locus in WTC11 dCK and the sgRNA donor plasmid used. E) FACS plot showing donor integration efficiency within landing pad at AAVS1 locus in WTC11 dCK PA01 PSCs. F) Graph showing repression efficiency of all promoter sgRNAs in Perturb-seq experiment in different cell lines and with different sgRNA delivery mechanisms. G) Heatmap showing correlation coefficient of promoter repression across cell lines and sgRNA delivery mechanisms. There is high correlation in promoter repression across all comparisons.

The disadvantages of lentiviral and PB-based sgRNA delivery mechanisms include the inability to control where sgRNAs integrate in the genome with potential epigenetic silencing during differentiation. Thus, we also tested sgRNA delivery with the PA01 large serine recombinase, which can mediate site-specific integration into the genome at 40-70% efficiency in K562 cells^17^. We first engineered WTC11-dCK-YS with a landing pad at the AAVS1 locus containing a PA01 attP sequence for site-specific integration and an expression cassette for PA01-GFP (**Figure 2D**). We designate these cells as WTC11-dCK-PA01. To test integration, we electroporated sgRNA-BFP donor plasmids and observed a recombination efficiency of ∼30% (**Figure 2E**), which is comparable to the rate of low MOI lentivirus infection. After enrichment for recombined cells by BFP+ cell sorting, we observed 90% knockdown of BEX3 expression (**Figure S2E**). Thus, WTC11-dCK-PA01 cells enable integration at a specific GSH site, consistent expression of sgRNA, and repression of specific targets.

Another advantage of recombinases is the ability to integrate tandem cassettes of U6-sgRNAs to test specific combinations of sgRNA in the same cell. As a proof of principle, we tested cassettes of 3 unique sgRNAs inserted into the AAVS1 locus of every cell. We wanted to assess 1) repression of target genes with multiple unique sgRNAs integrated into the AAVS1 locus and 2) position-dependent effects on repression. We found robust repression of target genes when the sgRNA was inserted in any of the 3 positions within the U6-sgRNA cassette (**Figure S2F(i-iii)**). The caveats for inserting multiple sgRNA cassettes into one locus are the lower integration efficiency of donor cassettes and the technical challenges of plasmid cloning in the presence of multiple U6-sgRNA cassettes.

To more systematically evaluate sgRNA delivery methods, we performed Perturb-seq experiments after CM differentiation. On average, the number of sgRNA integrations with each delivery method was comparable (average of sgRNA per cell: lentivirus=1.3; PB=1.4; recombinase = 1; **Supplementary Table 3**). However, we note that lentivirus delivery has more potential to controllably increase the number of integration events per cell by increasing titer. We observed robust and comparable levels of promoter and enhancer repression by either lentivirus or PB delivery of sgRNAs after differentiation to cardiomyocytes. This was true across PSC lines (WTC11-dCK and H9-dCK) and CRISPRi systems (H9-dCK and H9-idCK) (**Figure 2F-G, S2G-H**). For the WTC11-dCK lines, we note that Perturb-seq performance after sgRNA delivery by the PA01 recombinase was poorer compared to lentivirus and PB approaches (PA01 : 43% knockdown, lentivirus: 68%, PB: 67%, p-value = 9.23e-9, 1.02e-7 t-test) (**Figure 2F, S2G**). One possible reason for this poorer performance is the lower expression of sgRNAs integrated by PA01 at the AAVS1 locus. In summary, our results demonstrate a robust CRISPRi toolkit with efficient promoter and enhancer repression efficiency across cell lines and sgRNA delivery strategies, with potential uses in diverse Perturb-seq applications.

### Perturb-seq optimizations improve hit discovery and reduce cost

In addition to benchmarking pluripotent cell lines and sgRNA integration strategies, we also implemented several approaches to improve hit discovery and reduce cost **(Figure 3A)**:

- **<u>sgRNA design:</u>** While several computational methods can predict CRISPRi sgRNA efficiency, their performance has not been experimentally benchmarked. We evaluated the efficiency of several sgRNA design methods: Flashfry, CRISPR designer, and BLAST^18^. For each targeted promoter or enhancer, we designed up to 10 sgRNAs. We define ‘working sgRNA’ as a sgRNA that exhibits more than 50% of repression; about 70% of sgRNAs in all three methods are working sgRNAs. While we found no significant differences in knockdown efficiency between sgRNA design methods, testing multiple sgRNAs is required to identify hits (**Figure S3 A-B**).
- **<u>Maintaining sgRNA complexity</u>**: The number of cells sequenced per perturbation is a key determinant of a Perturb-seq experiment’s statistical power. Perturbations that alter cell proliferation will skew the sgRNA coverage of sequenced cells, with increasing impact over time. To maintain high sgRNA representation in Perturb-seq datasets, we developed a strategy to actively monitor representation by bulk sgRNA sequencing across PSC culture and differentiation using a rapid turnaround Nanopore sequencer (**Figure 3A,3C**). This critical quality control step enables the capability to abort scRNA-seq or adjust sequencing parameters depending on clonal expansion^19^.
- **<u>Cell sorting and superloading:</u>** To reduce the cost of Perturb-seq experiments, we optimized several parameters to maximize the yield of high-quality single cells sequenced (**Figure 3A**). First, we incorporated FACS sorting for BFP+ cells to increase the proportion of sgRNA+ cells sequenced. We sort for BFP+ cells after infection to enrich for sgRNA-containing cells (**Figure 3A, 3B**). Importantly, we sort for BFP+ cardiomyocytes before scRNA-seq library preparation to ensure only sgRNA-containing cells are sequenced (**Figure 3A, 3E**). Second, to ensure differentiation is technically progressing as expected we perform TNNT2 (marker for CM differentiation) FACS on day 8. We expect TNNT2 positive cells on day 8 to be at least 50% and on day 12 to be >90% (**Figure 3A, 3D**). Third, to increase the yield of sequenced cells, we use a cell hashing strategy coupled with super loading of cells (**Figure 3A, 3F, 3G**). We have found that loading ∼173k cells in one lane allows recovery of 60k high-quality single cells after quality filtering (see Methods; **Supplementary Table 3**). Importantly, we were able to significantly bring down costs incurred during Perturb-seq. To obtain 1,000 high quality cells for perturbing one enhancer/gene at MOI of 1, cost for library preparation and sequencing is ∼$48. Large-scale screens can decrease cost with higher MOI and lower cell number. Overall, our optimizations have helped us reduce cost and significantly increase the number of recovered transcriptomes after each perturbation.

**Figure 3:**
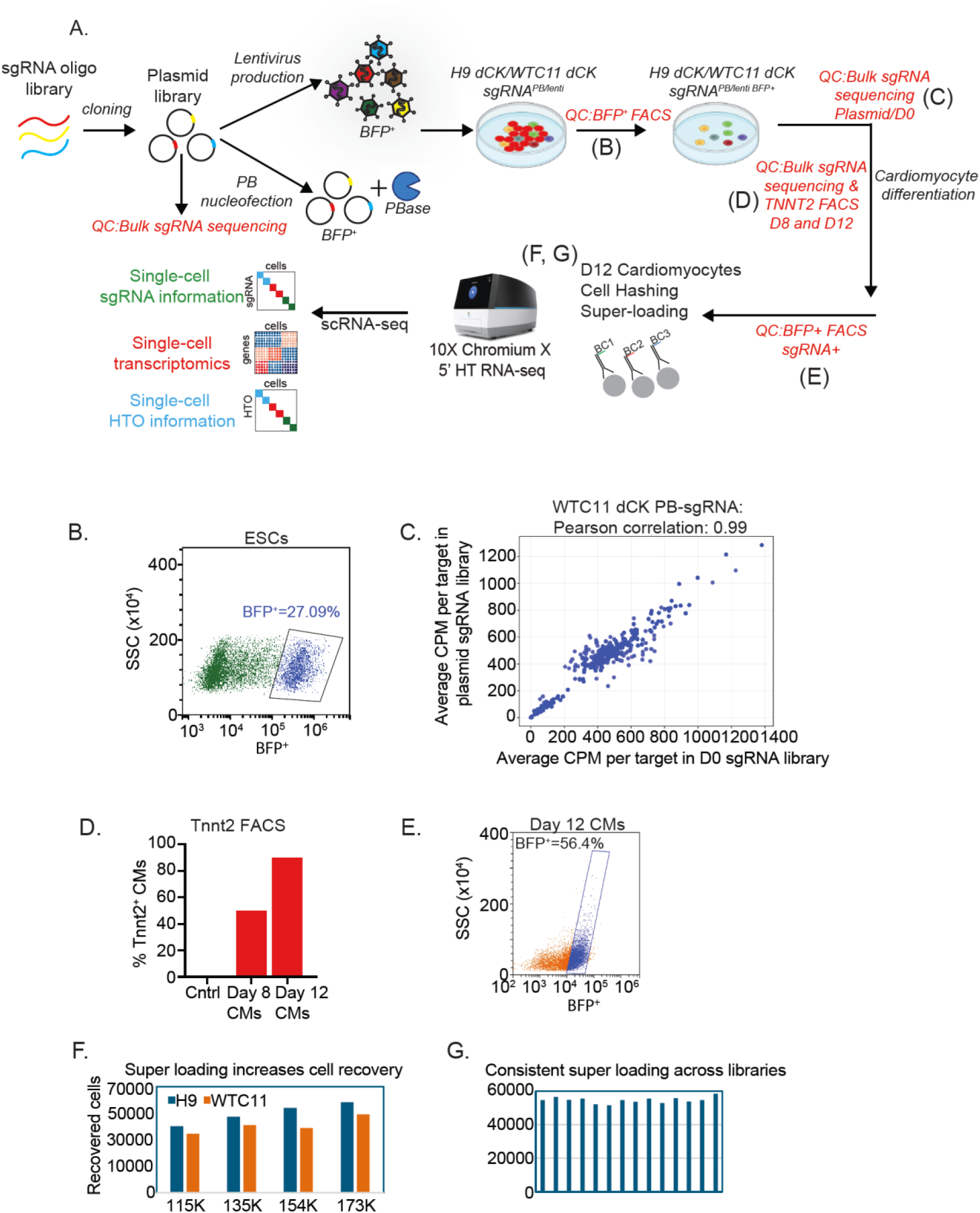
Perturb-seq workflow and key QC steps: A) Schematic of steps during Perturb-seq workflow including QC steps to ensure optimum sgRNA library coverage and efficient CM differentiation. B) FACS plot showing BFP positive PSCs selected after sgRNA integration. C) sgRNA sequencing of plasmid library correlates highly with sgRNA coverage in PSCs after integration. D) graph showing percentage of Tnnt2+ cells by FACS on day 8 and day 12 of CM differentiation. E) FACS plot showing BFP+ CMs were sorted and used for library preparation during scRNA-seq. F) Graph showing ∼60K cells were recovered after super loading during scRNA-seq. G) Graph showing consistent recovery of cells after super loading when preparing multiple libraries.

To facilitate rapid sharing of these optimized strategies for Perturb-seq in stem cells, we have assembled these standard operating procedures on Protocols.io^20^.

### Clonal expansion during Perturb-seq

Next, we tested if these engineered lines are compatible with Perturb-seq experiments in different lineages. We designed an sgRNA library targeting 64 transcription factors (TFs) expressed across lineages (**Figure S4A**) and performed Perturb-seq experiments in WTC11-dCK cells during cardiac and neural progenitor cells (NPC) differentiation. Overall, we observed strong transcriptional repression across iPSCs, CMs, and NPCs (**Figure S4B**). In all three lineages, we observe significantly increased recovery of cells with TP53 sgRNAs (**Figure 4A, 4C, S4C**), consistent with TP53’s role as a tumor suppressor. This was due to clonal expansion of p53-repressed PSCs, since plasmid library sequencing showed a more even distribution of sgRNAs (**Figure S4D**). The 6 sgRNAs targeting TP53 repressed TP53 mRNA levels by 90-99% (**Figure S4E**), indicating excellent CRISPRi silencing in CMs. Using a previously established H9 TP53^−/−^ PSC line, we confirmed that genetic deletion of TP53 leads to increased cell proliferation in cardiomyocytes **(Figure 4E**)^21^. Further, well-known cell cycle and apoptotic effectors of TP53 were dysregulated in TP53 knockdown PSCs and CMs as expected (**Figure 4B, 4D**). We further validated our single cell transcriptomic data by qPCR and observed that several cardiac specific genes were dysregulated in TP53 null PSCs and CMs compared to controls (**Figure 4F**). Thus, we implemented close monitoring of sgRNA complexity before and during differentiation to ensure robust representation of individual perturbations.

**Figure 4:**
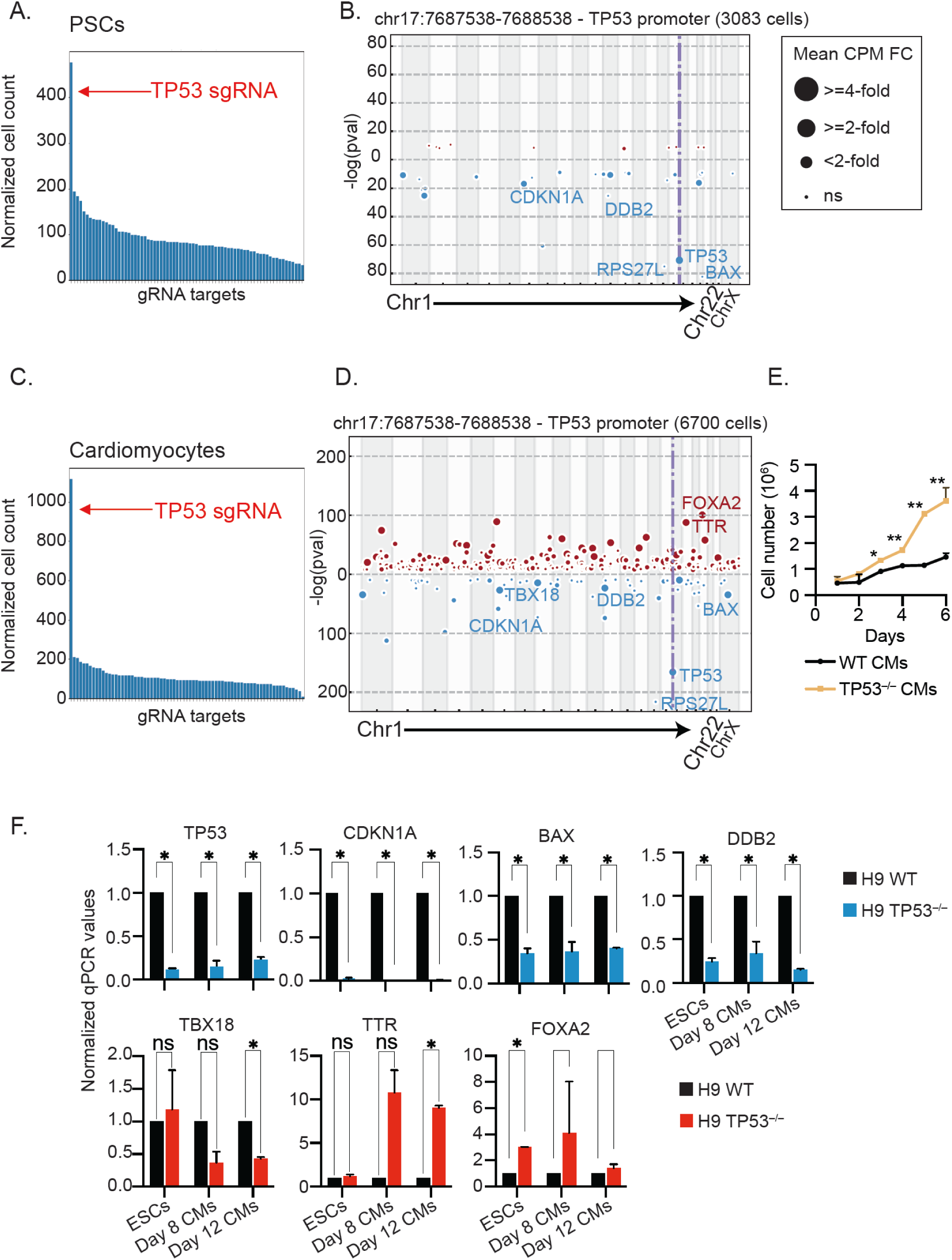
Clonal expansion during Perturb-seq: A) Bar plot showing number of cells with single sgRNA in PSC population. B) Manhattan plot showing up and downregulated genes upon repression of p53 in PSCs. C) Bar plot showing number of cells with single sgRNA in CM population. D) Manhattan plot depicting up and downregulated genes in CMs upon repression of p53.E) Growth curve of WT and TP53^−/−^ CMs. F) validation of scRNA-seq results in WT and TP53^−/−^ CMs by qPCR.

### Comparative analysis of regulatory networks across genes, enhancers, and stem cells

Our Perturb-seq experiments targeted several transcription factors (TFs), which orchestrate regulatory networks by controlling the expression of downstream genes. By measuring how TF knockdown causes global changes in gene expression, we sought to gain insights into how TFs functionally cluster by downstream regulatory effects. All results from our Perturb-seq experiments, including a list of perturbations and their effects on local and global gene expression, is provided in Supplementary Tables 4 and 5.

In WTC11 dCK^PB-sgRNA^ Perturb-seq, we observed significantly shared transcriptional effects for CRISPRi of NKX2-5, TBX20, and TBX5 (**Figure 5A, 5B**). Notably, perturbations of NKX2-5 or TBX20 result in overlap of 146 gene programs. Consistent with this, studies in mice have shown a direct cooperation between TBX20 and NKX2-5 to activate cardiac gene transcription during early development^22^. However, the correlation between TBX20 and TBX5 perturbations is lower (**Figure 5A-B**). In vertebrates, TBX20 and TBX5 have been shown to have distinct functions during heart development^23,24^. Additionally, repression of GRHL2 or SOX17 results in similar transcriptomes that are anti-correlated to perturbation of known cardiac TFs NKX2-5, TBX20, or TBX5 **(Figure 5A, 5C)**. This is consistent with SOX17 playing a key role in endoderm differentiation and implicates GRHL2 in similar developmental pathways^25^. We find very similar results in H9 dCK^PB-sgRNA^ Perturb-seq, indicating that the regulatory interactions are preserved in ESC- and iPSC-derived cardiomyocytes (**Figure 5C-D**).

**Figure 5:**
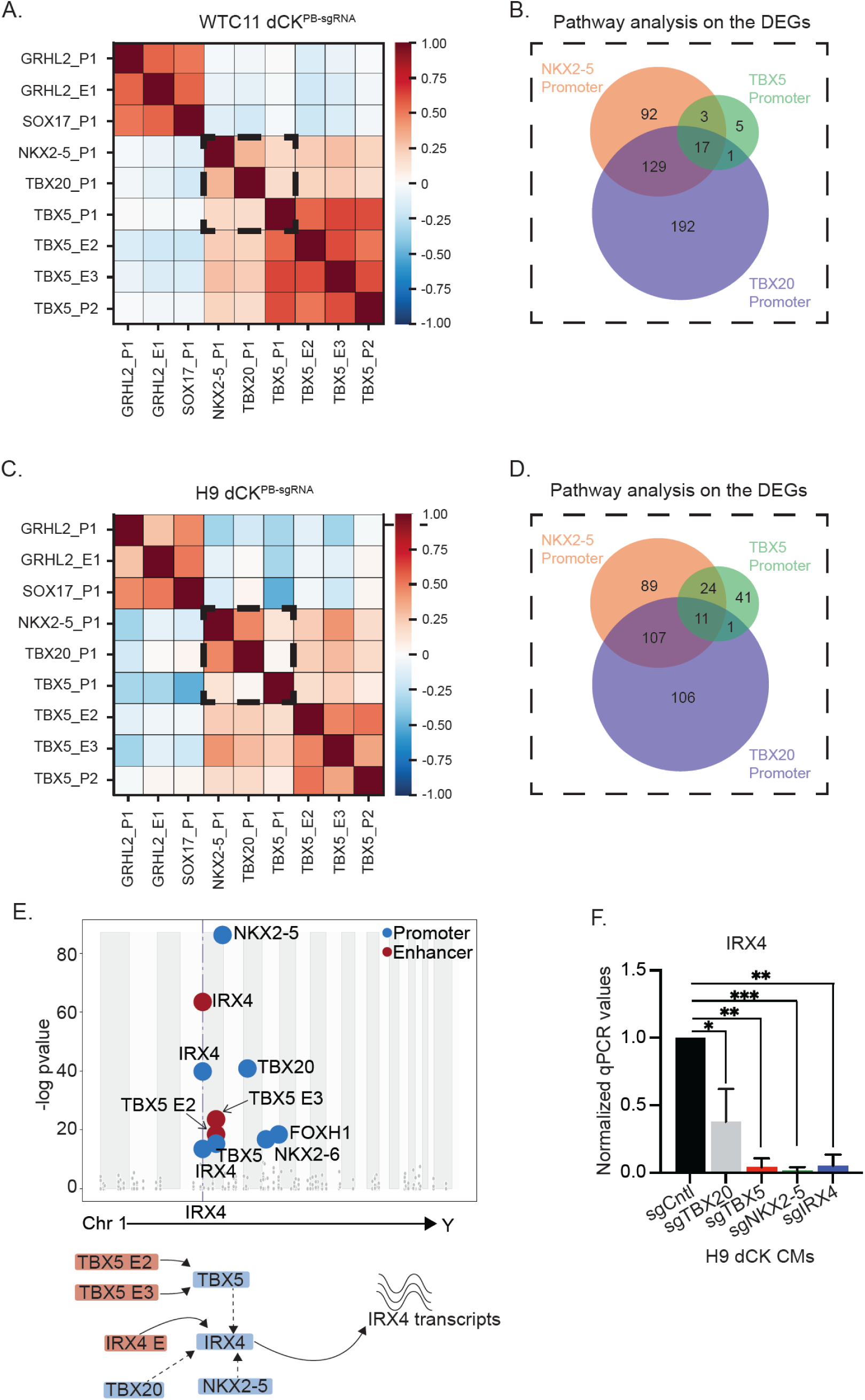
Constructing regulatory networks from Perturb-seq data: A) Correlation map showing groupings of perturbations in WTC11 dCK nucleofected with PB-sgRNA library. Groups are based on similarity between differentially regulated transcriptomes. Red squares indicate more similarity, and blue squares indicate poor correlation. B) Venn diagram showing number of biological pathways that overlap between perturbed CMs. C) Correlation map of differentially expressed transcriptomes in H9 dCK nucleofected with PB-sgRNA library. Red squares indicate high similarity between transcriptomes of indicated perturbation and blue squares indicate poor correlation. D) Venn diagram indicating biological pathways that overlap after each indicated perturbation. E) Perturbation of TBX5, TBX20 or NKX2-5 suppresses expression of IRX4 as indicated by scRNA-seq data. F) Validation by qpcr of IRX4 expression in CMs repressed for the indicated cardiac genes by CRISPRi sgRNAs.

Besides transcriptome-wide analysis, we also performed differential gene expression analysis, focusing on single regulatory loops that control CM development. For example, we find that CRISPRi of TBX5, NKX2-5, TBX20, GATA4, or NKX2-6 regulates expression of the well-known cardiac TF IRX4 (**Figure 5E**). Gratifyingly, this is consistent with prior mouse studies where the above cardiac TFs regulate IRX4 during heart ventricle development^26,27^. We validated these results by performing individual CRISPRi perturbations of TBX5, NKX2-5, and TBX20 and verifying that each individual gene repression resulted in a decrease in IRX4 expression by qPCR (**Figure 5F**). Thus, Perturb-seq enables the grouping of cardiac genes based on common downstream regulatory effects during development and discerns pathways that control CM differentiation.

In several instances, we were also able to extend this analysis to TFs and their corresponding enhancers. For example, in both H9 and WTC11 cells, we find that perturbation of the TBX5 promoter and its enhancers causes similar changes in the transcriptome (**Figure 5A, 5C**) and overlap of biological pathways (**Figure S5A-B**). Notably, CRISPRi of TBX5 enhancers and promoters converges on misregulation of the CHD-associated gene Versican (VCAN). Thus, our data positions TBX5 enhancers within a regulatory network that impacts proper cardiomyocyte differentiation (**Figure S5C**)^9^.

### Dissecting downstream effectors of NKX2-5 repression

Our TF Perturb-seq data also uncovered a novel regulatory connection where CRISPRi of NKX2-5 during cardiomyocyte differentiation leads to strong upregulation of NRG1 in both H9 and WTC11 CMs (**Figure 6A, S6A**). We observed that this regulation is specific to NKX2-5, as CRISPRi of TBX5 or IRX4 does not significantly upregulate expression of NRG1 (**Figure S6B**). Overall, there was a high overlap in gene programs upon NKX2-5 perturbation between H9 and WTC11 CMs (**Figure 6B, Figure S6C-D**).

**Figure 6:**
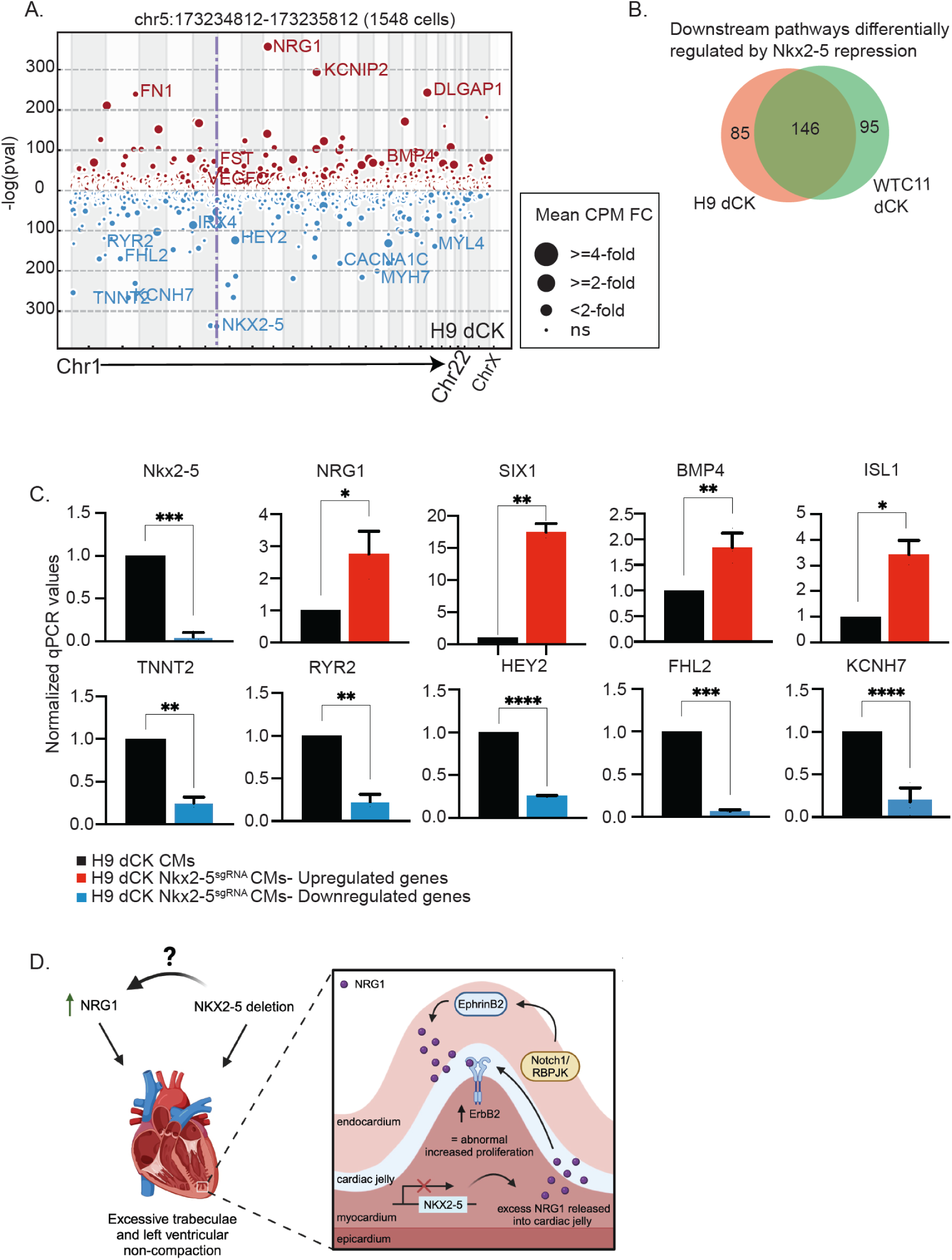
Identification of a novel NKX2-5 regulatory network in CMs. A) Manhattan plot showing differentially expressed genes upon NKX2-5 perturbation in H9 dCK CMs. B) Venn diagram showing high overlap in gene programs when NKX2-5 is repressed in H9 dCK and WTC11 dCk perturbed CMs. C) qPCR to validate scRNA-seq data in H9 dCK CMs repressed for NKX2-5 expression. D) Hypothesis on local regulatory loop between NKX2-5 and NRG1. From published literature NRG1 in endocardium regulates proper trabeculae formation in the myocardium. We have unraveled a novel loop in CMs where NKX2-5 repression upregulates NRG1 expression.

To validate these TF Perturb-seq results, we generated H9 NKX2-5 CRISPRi ESCs. We differentiated these cells into cardiomyocytes and confirmed altered expression of several cardiac genes, including ion channels (RYR2, KCNH7), sarcomeric proteins (TNNT2), signaling ligands (BMP4), and regulators of ventricle development (NKX2-5, HEY2, FHL2) (**Figure 6C**). There was a high concordance between our qPCR data and results obtained from Perturb-seq. For example, SIX1 is a transcription factor important for cardiac progenitor cell development and was reported to be upregulated in the hyper trabeculated hearts of NKX2-5^−/−^ mice^28,29^. Consistently, CRISPRi of NKX2-5 led to SIX1 induction in Perturb-seq and qPCR validation experiments (**Figure 6C**). A separate study showed that NKX2-5 directly represses expression of ISL1 to control cardiomyocyte differentiation^30^. In our Perturb-seq experiment, ISL1 was upregulated in NKX2-5 CRISPRi cells, and we confirmed its increased expression by qPCR (**Figure 6C**). Thus, we validated multiple downstream targets recovered after perturbation of NKX2-5.

Fetal heart development involves coordinated compaction of the inner trabecular layer of the ventricle with growth in the outer myocardial layer. NKX2-5 deletion during mouse embryonic development results in a ventricular hyper-trabeculated phenotype^29^. Both Notch1 signaling and Neuregulin-1 (NRG1) activity have been shown to be required for proper trabeculation, particularly through endocardium-myocardium communication^31–33^. Endocardial NRG1 has been shown to bind myocardial ERBB4 to promote downstream signaling that results in cell growth and migration^34^. Consistent with this, mice lacking NRG1, ERBB4 and its coreceptor ERBB2, have defects in ventricular trabeculation and die in utero^35–38^. Taken together, our results provide experimental support for the hypothesis that NKX2-5 represses NRG1 expression in cardiomyocytes to ensure proper trabeculation and ventricle development (**Figure 6D**).

## DISCUSSION

Perturb-seq is a powerful approach to understand the impact of genes and regulatory elements in a dynamic stem cell model of human development. However, these studies can be technically challenging and expensive. In this study, we optimized several parameters for the systematic application of Perturb-seq in differentiated PSCs. We are using these standard operating procedures to scale Perturb-seq experiments in cardiomyocytes and neurons, as part of the IGVF Consortium. The detailed procedures can be found on Protocols.io^20^. The engineered stem cell lines, sgRNA plasmids, and other reagents are being deposited to open repositories for use by the scientific community. Below, we discuss several key parameters:

- **<u>dCas9-KRAB expression:</u>** Stable expression of dCK from a genomic safe harbor is essential to ensure consistent expression of the effector and optimum repression of sgRNA target regions. Previously, we used PiggyBac and lentiviral integration of dCK into the PSC genome, which led to variegated expression and silencing of the vector over time. To obtain clonal, non-variegated expression of the dCas9 effector, we engineered several stem cell lines (H9 ESC and WTC11 iPSC) in which dCK was inserted into the CLYBL locus, similar to previous studies in neuronal models^13,14^. We also opted to use heterozygous cell lines over homozygous dCK cell lines in our Perturb-seq screens during CM development for three reasons. First, one copy of the WT allele was left untouched for further engineering. Second, we wanted to avoid potential detrimental effects if the locus has an unknown function during CM development. Third, dCas9-KRAB expression is not limiting for efficient target gene repression (data not shown).
- **<u>sgRNA delivery:</u>** Our optimizations indicate that sgRNA expression is a critical parameter for Perturb-seq. Although not discussed in depth, we found that placing the sgRNA cassette (U6-sgRNA) upstream of the selection cassette (EF1a-puro-tagBFP) was crucial, suggesting that Pol II read-through may negatively impact performance. These results also suggest that sgRNA expression is more limiting than dCK expression. This could be one reason that sgRNA integration by recombinase performed worse than lentiviral and PB approaches, as the AAVS1 landing pad is subject to read-through by the endogenous AAVS1 gene. Adopting other safe harbors for sgRNA integration could improve sgRNA expression and target repression. Separately, while this study performed Perturb-seq with low MOI (1 sgRNA/cell), the ability to control the MOI of sgRNA infection is a critical parameter to economically scale Perturb-seq experiments so that each cell can provide information when multiple targets are repressed using several sgRNAs.
- **<u>Quality control steps:</u>** Perturb-seq experiments in stem cell models can take several weeks, which incurs the high cost of culture and differentiation of stem cells in large batches (the CM experiments take >30 days from start to finish). Problems like poor differentiation and loss of sgRNA complexity will negatively impact the quality of resulting datasets and the insights that can be gained. To combat this issue, we have developed steps to actively monitor data quality across the experiment: measuring TNNT2 by FACS to ensure efficient differentiation, performing BFP sorting on cells to enrich for cells with sgRNAs, and examining sgRNA complexity over time. These steps help to maximize Perturb-seq data quality and minimize cost.

The Perturb-seq datasets generated in our differentiation systems are highly consistent, even across experimental parameters, such as stem cell line and sgRNA delivery approach. These observations are indicative of robust repression and high data quality. Therefore, we used our datasets to define regulatory interactions between well-known cardiac transcription factors and to identify regulatory networks linking enhancers to the TFs they activate. We used Perturb-seq to highlight a potentially new role for NKX2-5 in ventricular formation. We found that repression of NKX2-5 leads to upregulation of NRG1, suggesting that active repression of NRG1 by NKX2-5 in CMs allows for proper trabeculation and hence heart ventricle formation during development. Thus, our Perturb-seq results unveiled a novel regulatory network that may impact CM differentiation.

In summary, we have established a working pipeline that will be used to test the function of several promoters and enhancers during differentiation. Further, it allows us to test effects of variants on development and disease progression. Thus, we foresee our tools will benefit the scientific community and aid others in performing similar CRISPRi experiments in biomedically-relevant iPSC-derived models.

## SUPPLEMENTARY FIGURES

**Figure S1:**
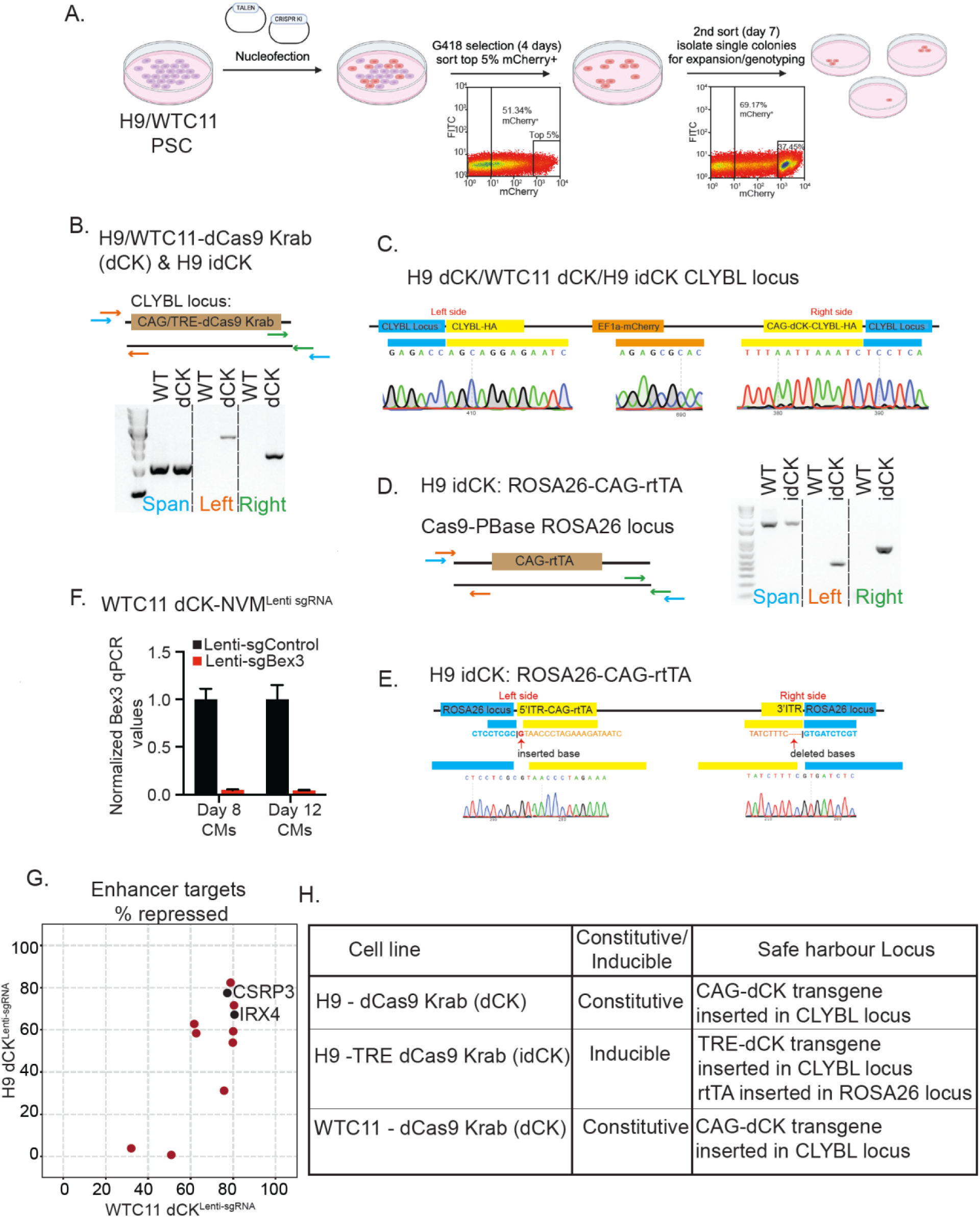
Details of GSH integration and testing CRISPRi repression. A) Cartoon showing steps followed to engineer dCK PSCs. FACS based sorting for EF1-mCherry was performed twice sequentially to enrich for pure population of mcherry positive dCK PSCs. B) PCR genotyping of CLYBL locus in H9/ WTC11/H9 idCK PSCs shows correct integration of transgene in one allele. The arrows in the locus diagram indicate location of PCR primers. C) Sequencing traces showing correct integration of transgene at CLYBL locus. D) Genotyping strategy to PCR the CAG rtTA transgene at ROSA26 GSH locus in the H9 idCK line. The arrows indicate location of the PCR primers within the locus. E) sequences traces showing correct integration of CAG-rtTA at ROSA26 locus in H9 idCK PSCs. F) qPCR showing Bex3 repression by lentiviral sgRNAs in WTC11 dCK NVM PSCs. G) Correlation of enhancer repression across H9 dCK and WTC11 dCK infected with lentiviral sgRNAs. CSRP3 and IRX4 enhancers are repressed in both H9 dCK and WTC11 dCK PSCs. H) Table detailing the PSCs engineered in both H9 and WTC11

**Figure S2:**
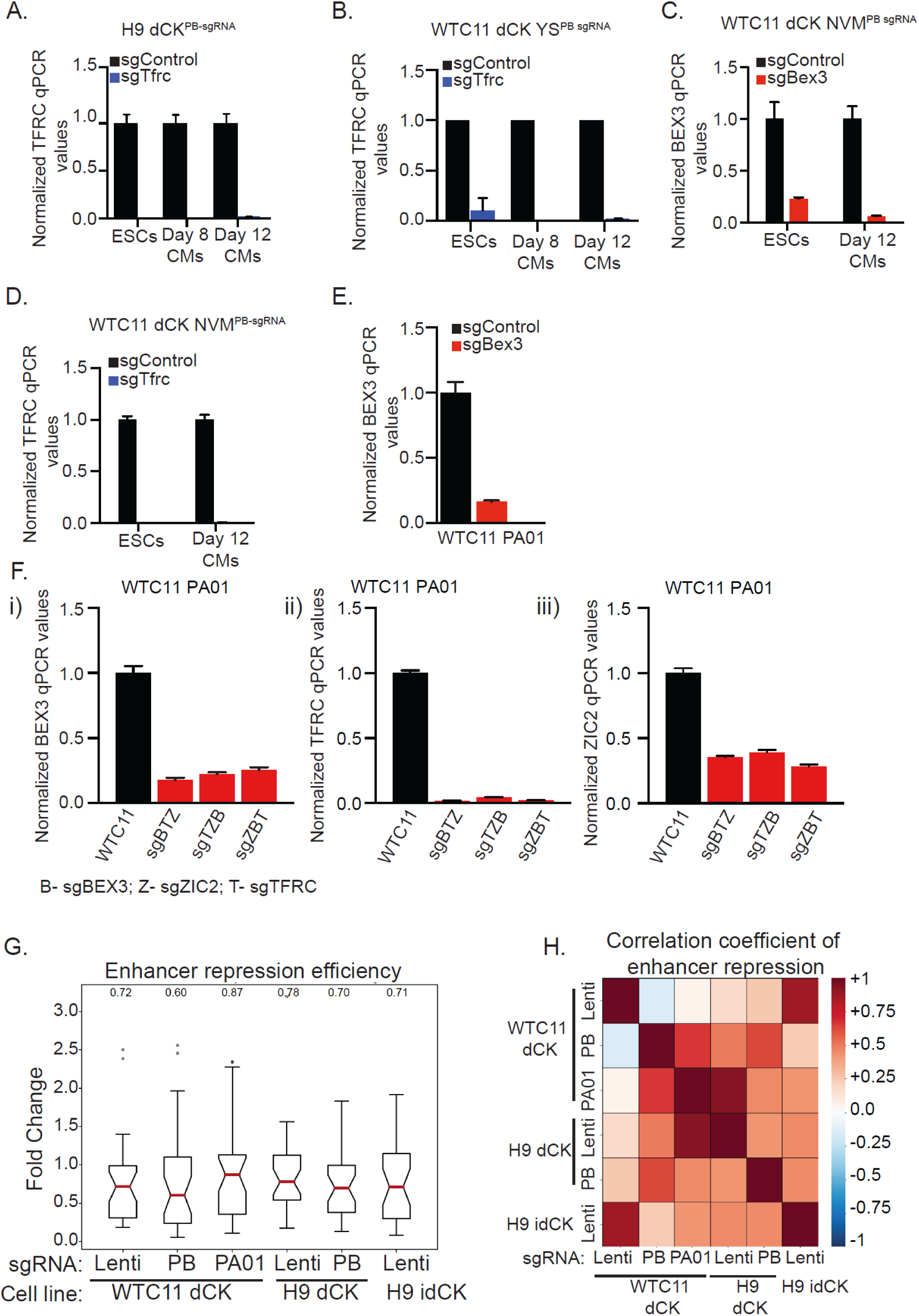
Impact of sgRNA delivery method on target repression. A) Graph showing repression of TFRC in H9 dCK ESCs and CMs using PB-sgRNAs by qPCR. B) Graph showing repression of TFRC in WTC11 dCK YS ESCs and CMs using PB-sgRNAs by qPCR. C&D) Graph showing repression of BEX3 (C)or TFRC (D) in WTC11 dCK NVM ESCs and CMs using PB-sgRNAs by qPCR. E) Graph showing repression of BEX3 in WTC11 dCK PA01 ESCs by qPCR. F) Graph showing repression of Bex3 (i), TFRC (ii), ZIC2 (iii) in PA01 line using cassettes of three sgRNAs. Position of sgRNA within the cassette doesn’t inhibit repression of targets. G) Graph showing repression efficiency of all enhancer sgRNAs in Perturb-seq experiment in different cell lines and with different sgRNA delivery mechanisms. H) Heatmap showing correlation coefficient of enhancer repression across all cell lines and delivery mechanisms. There is a low correlation in enhancer repression across all comparisons.

**Figure S3:**
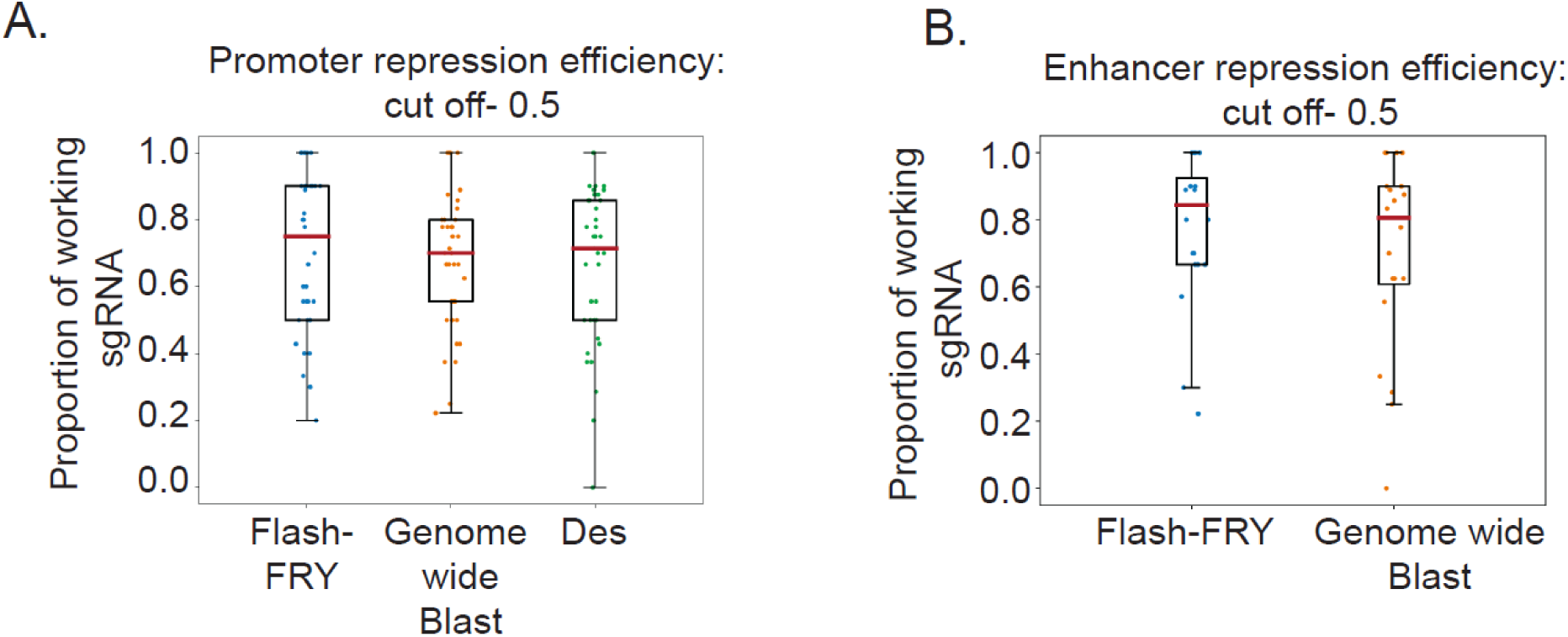
Impact of sgRNA design method on target repression: (A) The box plot shows that the proportions of working sgRNAs for promoters are similar across different computational sgRNA design methods. The working sgRNA is defined as at least 50% repression level, the proportion is calculated with individual perturbation region: (# of working sgRNA) / (# of all sgRNA). (B)The box plot of working sgRNAs for enhancers are similar across computational sgRNA design methods.

**Figure S4:**
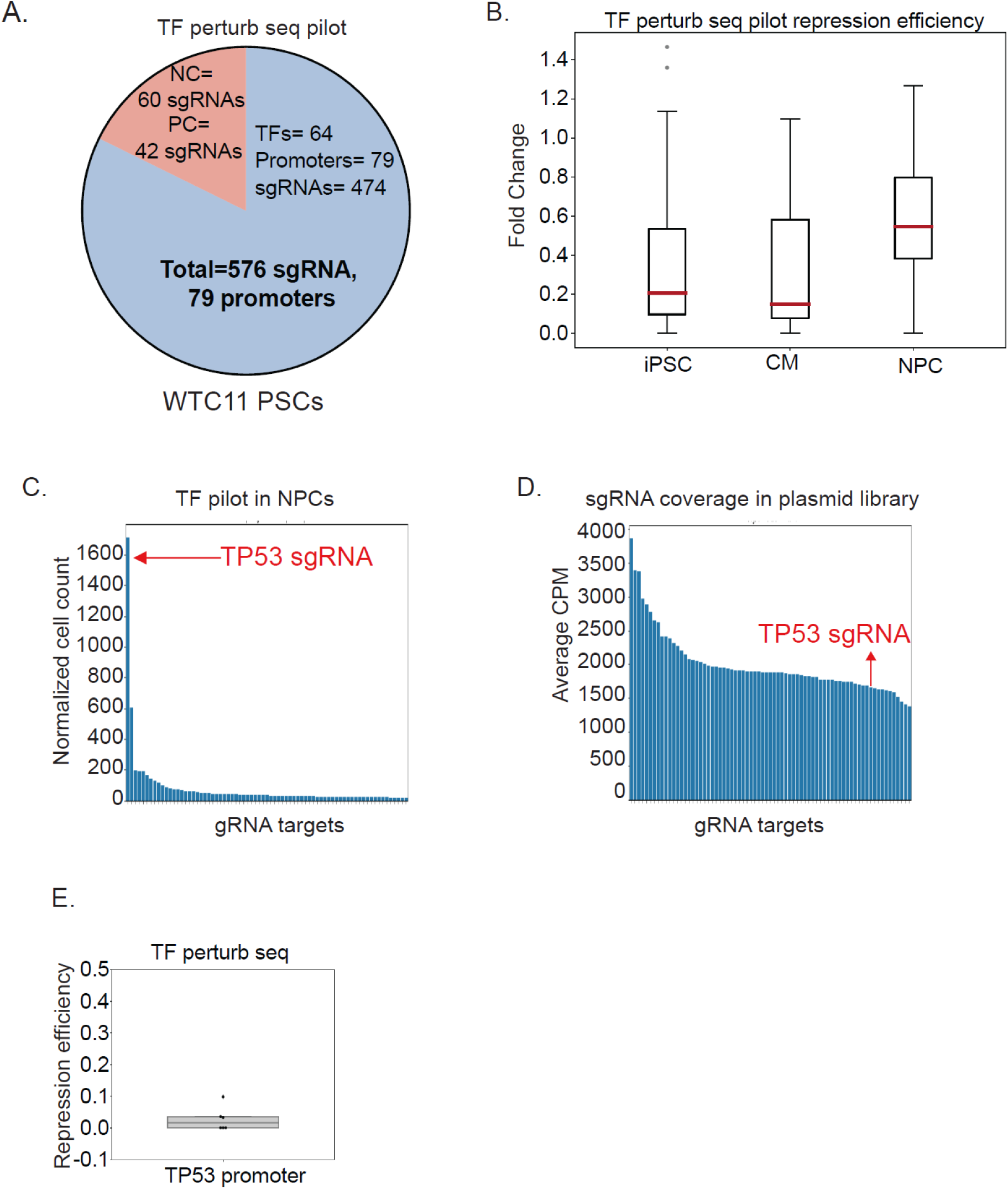
TF Perturb-seq pilot parameters and NPC clonal expansion: A) Venn diagram showing distribution of sgRNA in TF pilot Perturb-seq experiment. B) Repression efficiency of all sgRNAs in iPSCs and differentiated CMs or NPCs during Perturb-seq. C) Bar plot showing number of cells with single sgRNA in NPC population. D) Bar plot showing coverage of sgRNAs in plasmid library. E) Graph showing repression efficiency of p53 promoter in CMs in TF pilot Perturb-seq experiment. Each dot represents a single sgRNA.

**Figure S5:**
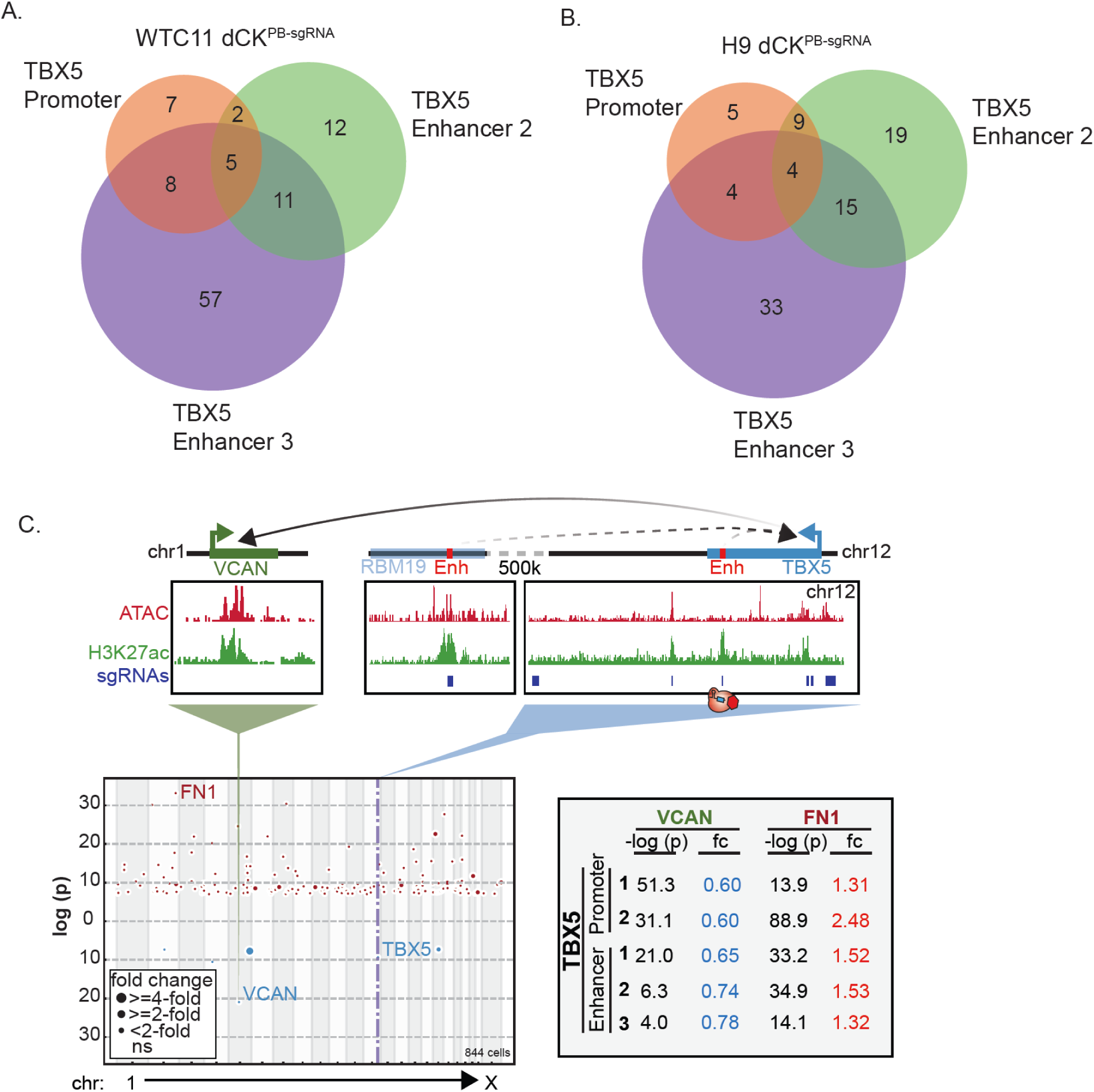
Perturb-seq allows us to group targets based on differentially expressed transcriptomes: (A) Venn diagram showing number of biological pathways that overlap between TBX5 promoter and enhancer perturbed WTC11 dCK CMs. (B) Venn diagram showing number of biological pathways that overlap between TBX5 promoter and enhancer perturbed WTC11 dCK CMs. (C) TBX5 promoters and enhancers regulate expression of the CHD associated gene VCAN. Manhattan plot shows the differentially regulated genes upon TBX5 repression. Blue dots indicate downregulated genes including TBX5 and VCAN and red dots indicate upregulated genes including FN1.

**Figure S6:**
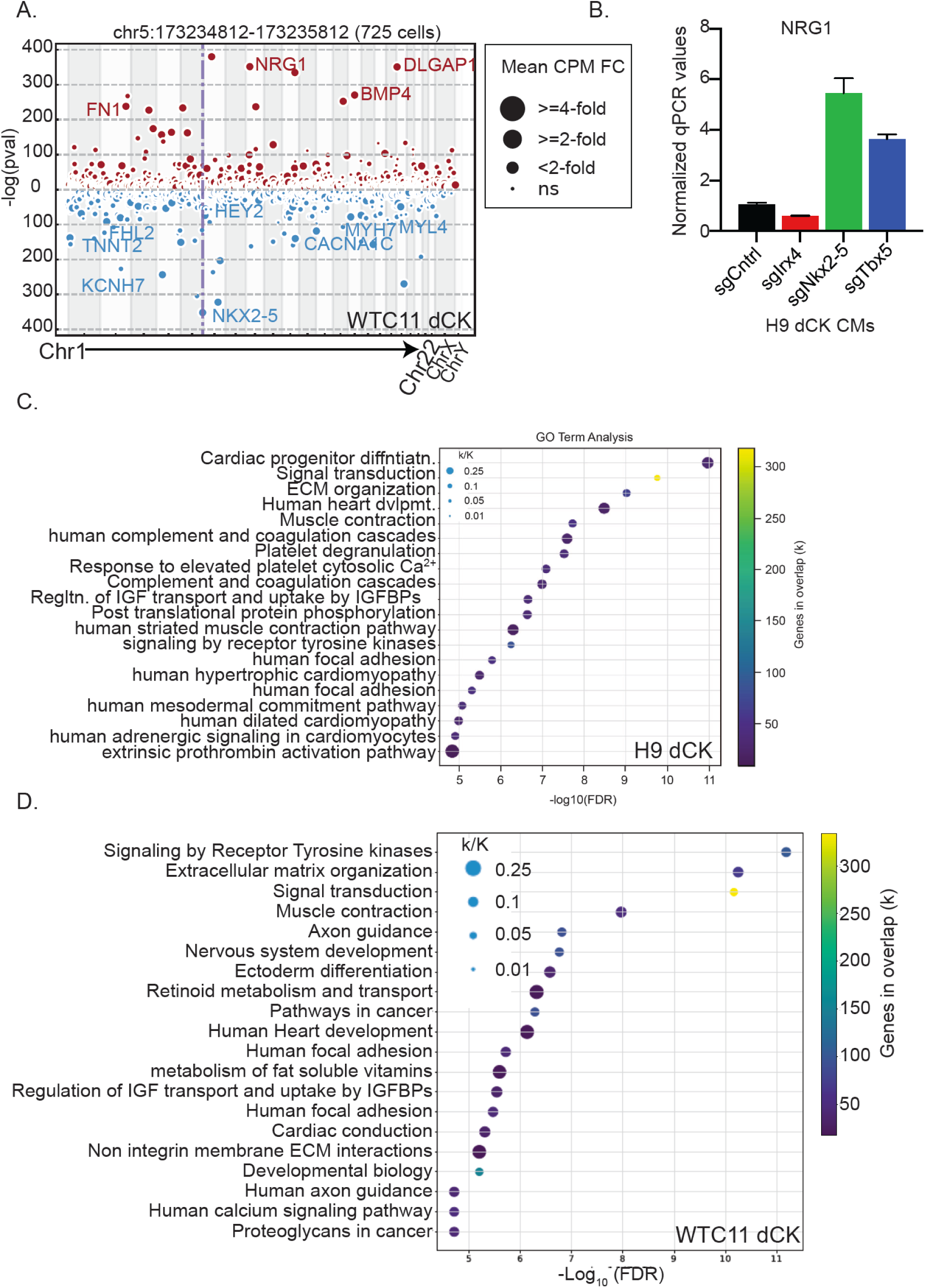
Effect of NKX2-5 repression on CM differentiation and NRG1 expression: (A) Manhattan plot showing differentially expressed genes upon NKX2-5 perturbation in WTC11 dCK CMs. B) Graph showing upregulation of NRG1 in NKX2-5 repressed CMs by qPCR. C) GO term enrichment analysis on differentially expressed genes upon NKX2-5 perturbation in H9 dCK CMs. (D) GO pathway analysis on differentially expressed genes upon NKX2-5 perturbation in WTC11 dCK CMs.

## SUPPLEMENTARY TABLES

**Supplementary Table 1:** Comparison of sgRNA delivery mechanisms.

**Supplementary Table 2:** Validation oligo sequence / qPCR primer sequences.

**Supplementary Table 3:** Library QC stats.

**Supplementary Table 4:** Differential expressed genes results (local analysis).

**Supplementary Table 5:** Differential expressed genes results (global analysis)

## METHODS

### Engineering of H9 dCK, H9 idCK, WTC11 dCK ESCs

Human pluripotent stem cells were grown in mTeSR plus and passaged using PBS+EDTA every 3-4 days. For nucleofection, PSCs were fed with CloneR2/mTeSR Plus 3 hours before dissociation with 1X TrypLE. Two million cells were resuspended in 100 ul P3 buffer (Lonza Inc.) and mixed with plasmid DNA, transferred to cuvette, and pulsed using the CB156 program. The cells are immediately diluted in 500 ul CloneR2/mTeSR Plus, incubated at 37 C for 10 minutes and plated in CloneR2/mTeSR Plus. Three days post nucleofection positive cells were single cell sorted and plated under selection. Cells were again sorted and plated at low density under selection and antibiotic-resistant clones were isolated and genotyped. To engineer H9/WTC11 dCK lines, 10 ug of CAG-dCas9-KRAB donor and 5 ug of each CLYBL TALEN vectors were nucleofected into cells, sorted for mCherry and then selected with 100ug/ml G418. To engineer H9 idCK PSC, first rtTA was inserted into ROSA26 locus by nucleofecting 1.45 ug hypaCas9-PBase(-)Int, CAG-rtTA donor and TRE-EGFP reporter (molar ratio 1:2.5:2.5). Cells were sorted for EGFP, selected with 100ug/ml G418 and then TRE-dCas9 KRAB was inserted in CLYBL locus using TALENS. 10 ug of CLYBL TRE-dCas9-KRAB donor, 5 ug of each CLYBL TALEN vectors was nucleofected into PSCs, sorted for mCherry and selected with 0.25 ug/ml Puromycin. Single cell clones were genotyped to verify correct integration of transgenes.

### Genomic DNA extraction and genotyping

Genomic DNA was extracted from H9 ESCs using Qiagen DNeasy Blood & Tissue Kit (Catalog #69504). Plasmid Integration was confirmed via PCR using APEX 2X Taq PCR Master Mix (Catalog #42-138B) according to manufacturer’s instructions.

### Nucleofection of sgRNAs into H9/WTC11 dCK ESCs

H9 ESCs were nucleofected using 10ug piggybac U6-sgRNA-puro plasmid using NEPA nucleofector. Lentiviral infection of sgRNAs was done as previously described ^9^. The cells were allowed to recover in mTESR plus CloneR2 for 72 hours. Puromycin selection was applied at 1ug/ml for 7-10 days. PSCs were differentiated into CMs or NPCs according to our established protocol and qPCR was done to validate results from Perturb-seq ^9,21^.

### qPCR to validate Perturb-seq data

RNA was extracted from differentiated H9 ESCs using Qiagen RNeasy Mini Kit (Catalog #74104) and single-stranded cDNA was synthesized using Lunascript RT SuperMix Kit (Catalog #M3010X). To quantify the target transcript levels, qPCR was performed with Med Chem Express Sybr Green Master Mix (Catalog #HY-K0523-2000) using the CFX384 Real Time PCR System (Bio-Rad).

### sgRNA enrichment/dropout

To analyze the clonal expansion of certain gRNAs in the TF perturb seq experiment. We first identified all cells that had both transcriptome and gRNA information. Then, we created a binary table of gRNA in a cell. Then, gRNAs targeting the same regions were merged together. Finally, the number of cells per targeted region were counted. We normalized the cell count to their frequency in the plasmid library.

### Perturb-seq

The full Perturb-seq protocol can be found on Protocols.io ^20^.

### Next generation sequencing

All the libraries are sequenced with the Illumina platforms, the details of sequencing parameters can be found on the IGVF data portal.

### Analysis details

- **Perturbation region selection** Non-coding regulatory elements are defined by the enrichment of ATAC-seq peaks and H3K27ac ChIP-seq peaks ^39,40^. Public available datasets are downloaded with sra_toolkit (version=2.8.2-1), and mapped with BWA MEM (version=0.7.5). The peaks are called with macs2 (version=2.1.2). The ATAC-seq peaks of all the time points are merged, and extended +/- 250bp from the peak center. H3K27ac ChIP-seq signals are quantified within the extended open chromatin region, and normalized with RPKM (Reads Per Kilobase per Million mapped reads). The cutoff of H3K27ac signal (log2 RPKM) is set to be greater than 1.5. Cardiac genes are selected based on our previous publication and IGVF cardiac metabolic focus group ^9,12^.
- sgRNA design For the Pilot Pilot library, the sgRNAs are selected from three different algorithms: Flash Fry, CRISPR Designer and Genome-wide blast. Around 10 sgRNAs are selected for each method. For the TF Perturb-seq pilot library, 6 sgRNAs are selected from Horlbeck et al for each TF promoter ^41^. Perturb-seq QC Single cell transcriptome libraries are mapped to the human reference genome (hg38) using Cell Ranger (version=7.0.0). CellPlex libraries and sgRNA libraries are mapped with FBA (version=0.0.14)^42^. Both ambient CellPlex and sgRNA reads are filtered with the saturation curve method described by Drop-seq. The analyzed cells of all datasets in this study are 1 CellPlex tagging (experimental singlet) and more than 1 sgRNA per cell. The detailed information for each dataset is listed in **Supplementary Table 3**.
- Differential expression (DE) analysis pySpade (version=0.1.2) is used for differential expression analysis^43^. To generate the files for downstream analysis, pySpade “process” function is applied in the Cell Ranger output file to generate transcriptome and sgrna matrices (h5 files). pySpade “fc” is used to ensure the positive controls exhibit strong repression in each experiment. Complementary cells are used for the background. “DEobs” and “DErand” are utilized to perform global DE analysis. The randomization method parameter of “DErand” is set to be “equal”. The bins for the number of cells are included in **Supplementary Table 4&5**. Manhattan plots are generated with pySpade “manhattan” with parameters reflecting FDR < 0.1.
- Global DE genes correlation To examine whether perturbing promoters and enhancers of the same gene exhibit similar transcriptome alteration, we perform the pairwise correlation analysis of all the perturbation regions within the same screen. Significant DE genes are considered in the analysis. The p-values of both up-regulated and down-regulated genes are set for different signs (positive value or negative value). Pearson correlation analysis is applied in the pairwise comparisons. To visualize the similarity, hierarchy clustering is used to group the regions.
- Pathway comparison To compare the enriched biological pathways for each perturbation, differential expressed genes are the input of gseapy (version=0.14.0) “enrichr” function, and expressed genes (genes expressed in at least 5% of cells in the single cell libraries) are used as background^44^. Several gene sets are compared including ‘BioCarta_2016’, ‘KEGG_2021_Human’, ‘Reactome_2022’, and ‘WikiPathways_2019_Human’. The adjusted p-value cutoff is 0.05.

## DATA AVAILABILITY

The complete FASTQ data, sgRNA sequences, Differential Expression tables and all relevant metadata is available on the IGVF Consortium Data Portal (https://data.igvf.org/). The data used throughout this paper can be accessed through the following accession numbers and links:

- /analysis-sets/IG CFDS 9192 YMGI/
- /analysis-sets/IGVFDS6924DJAZ/
- /analysis-sets/IGVFDS6332VCTO/
- /analysis-sets/IGVFDS4003HZAB/
- /analysis-sets/IGVFDS5498SDRJ/
- /analysis-sets/IGVFDS7340YDHF/
- /analysis-sets/IGVFDS2266YDVM/
- /analysis-sets/IGVFDS2873EPIU/
- /analysis-sets/IGVFDS4252LFRU/
- /analysis-sets/IGVFDS4762GRIW/
- /analysis-sets/IGVFDS4389OUWU/

## CODE AVAILABILITY

The analysis scripts are available on GitHub (https://github.com/Hon-lab/ESC_engineering).

## ACKNOWLEDGEMENTS

We acknowledge the BioHPC computational infrastructure at UT Southwestern for providing HPC and storage resources that have contributed to the research results reported within this paper. G.C.H is supported by CPRIT (RP190451), NIH (DP2GM128203, UM1HG011996), the Burroughs Wellcome Fund (1019804), and the Green Center for Reproductive Biology. N.V.M. was supported by the NIH (HL136604, HL151650, HG012768, and HG011996), the Burroughs Wellcome Fund (1009838), and the Department of Defense (PR172060). We thank members of the Hon and Munshi laboratories at UTSW for critical suggestions and members of the IGVF consortium for methodological feedback. We thank Yin Shen (UC-San Francisco) for providing the WTC11-dCK cell line.

